# Tailor-made sRNAs: a toolbox to control metabolic targets

**DOI:** 10.1101/801027

**Authors:** Patrícia Apura, Alexandra Peregrina, Margarida Saramago, Sandra C. Viegas, Sandra M. Carvalho, Lígia M. Saraiva, Susana Domingues, Cecília M. Arraiano

**Affiliations:** Control of Gene Expression Lab, Instituto de Tecnologia Química e Biológica António Xavier, Universidade Nova de Lisboa, Oeiras, Portugal; Molecular Mechanisms of Pathogen Resistance Lab, Instituto de Tecnologia Química e Biológica António Xavier, Universidade Nova de Lisboa, Oeiras, Portugal

**Keywords:** Synthetic biology, control of gene expression, small non-coding RNAs, Hfq, *Pseudomonas putida*

## Abstract

*Pseudomonas putida* is a highly attractive production system for industrial needs. Modulation of gene expression is an urgent need to redesign *P. putida* metabolism for its improvement as biocatalyst at industrial level. We report the construction of a small RNA-based system with potential to be used for different purposes in synthetic biology. Due to their modular composition, design facilities and ability in tuning gene expression, sRNAs constitute a powerful tool in genetic and metabolic engineering. In the toolbox presented here, the synthetic sRNA is specifically directed to any region of a chosen target. The expression of the synthetic sRNAs is shown to differentially modulate the level of endogenous and reporter genes. The antisense interaction of the sRNA with the mRNA results in different outcomes. Depending on the particularity of each sRNA-target mRNA pair, we managed to demonstrate the duality of this system, able either to repress or overexpress a given gene. This system combines a high specificity with a wide applicability due to its ability to modulate the expression of virtually any given gene. By plugging-in and -out genetic circuits, this tailor-made regulatory system can be used to redesign *P. putida* metabolism, fulfilling an important industrial gap in synthetic biology.

## Introduction

Modulation of protein expression has been a challenging task (Mahalik *et al.* 2014). The main problems arise from the accumulation of toxic intermediates, titration of co-factors and overburdening of the cell factory, which compromise cellular productivity. Some problems can be overcome with an adjustment of enzymes’ levels using a combinatorial expression library (Lee *et al.* 2013). Typically, this implies the creation of libraries with changes in the involved genes. An ideal system would not rely in pre-constructed libraries, would have a dynamic range, could be programmable and would be non-toxic for the host. Good candidates for such system are small noncoding RNAs (sRNAs) (Ghodasara *et al.* 2017).

sRNAs are functional RNA molecules usually not translated and involved in many different cellular functions (Hor *et al.* 2018). Their versatility is attributable to their capacity to interact with RNA, DNA, proteins and small molecules. Most sRNAs directly base-pair with mRNA(s) to regulate one or more targets. *Cis*-encoded sRNAs are located on the opposite strand of their targets, having perfect complementarity. *Trans*-encoded sRNAs have a different genomic location and partial complementarity. Usually, the RNA chaperone Hfq helps binding to the targets and protects sRNAs from degradation (Wagner *et al.* 2015). Stretches of RNA located in the 5’- or 3’-end of mRNAs can function as sRNAs mediating RNA conformational changes. These are called riboswitches, and constitute the majority of synthetic sRNAs developed so far. They typically contain a sensor domain, plus a regulatory domain. Most require the fusion of target sequences to natural or modified sRNA scaffolds, which implies that they must be designed *ad hoc* for each target (Amman *et al.* 2012).

Inspired by the enormous versatility of RNA regulators, in this work we have explored *trans*-acting sRNAs since they allow the fine-tuning of desired genes and the identification of essential genes which cannot be identified by gene-knockout experiments (Yoo *et al.* 2013). Most studies have relied on random screening, but Na *et al.* showed the application of synthetic sRNAs rationally designed (Na *et al.* 2013). Based in this synthetic sRNA system developed for *Escherichia coli*, we have developed a new gene modulation system in *Pseudomonas putida*, considered one of the best hosts for optimization of product formation at the industrial level (Nikel *et al.* 2014). *P. putida* can withstand much stress, having higher productivity than other bacteria on a large scale. With this work we hope to have contributed to fill some of the gaps on industrial yields.

Our synthetic sRNAs comprise two parts: a seed sequence, which is a complementary sequence to the target mRNA (it can be directed to any region of the target), and a scaffold sequence that is responsible for the recruitment of the RNA chaperone, Hfq. The interaction of the sRNA with the mRNA is expected to result in different outcomes, such as repression or improvement of target mRNAs translation, by changing access of the ribosomes to the Ribosome-binding site (RBS), and/or protection of mRNA transcript from ribonuclease cleavage. Indeed, we managed to differentially modulate the expression of endogenous and reporter genes.

The largest advantage of the system here presented is that it is a dual-system that can be applied to either increase or decrease the amount of protein produced by simply directing the synthetic sRNA to different regions of the target mRNA (5’-end, coding region or 3’-end). As such, it can be used to redesign the metabolism of *P. putida* improving its potential for biocatalysis at the industrial level.

## Experimental Procedures

### Bacterial Strains, Plasmids and Growth conditions

Strains and plasmids are listed in Table S1, and oligonucleotides in Table S2. Strains were grown in Luria-Bertani (LB) supplemented with the antibiotics kanamycin (50 µg/ml), gentamicin (10 µg/ml) or carbenicillin (100 µg/ml), whenever required. *E. coli* was grown at 37°C and *P. putida* at 30°C, both under agitation (180 rpm).

pSEVA238 (Silva-Rocha *et al.* 2013) was amplified with primers containing a 5’-tail with the MicC scaffold sequence (Na *et al.* 2013). Primers SMD154 and SMD155 contain respectively the upstream and downstream region of the MicC scaffold. The 3’-end of these primers is complementary to pSEVA238 so that the MicC scaffold could be inserted exactly at the +1 of the XylS/*Pm* promoter. The amplified fragment was purified with NucleoSpin Gel and PCR clean-up (Macherey-Nagel) and ligated with T4 DNA Ligase (Thermo-Fisher). This construction (named pSEVA238-MicC) was transformed into KT2440 wild-type, KT-GFP and KT-YFP strains, used as controls in fluorescence assays.

To insert seed sequences (20-25 nts) into pSEVA238-MicC upstream of MicC scaffold, the same PCR amplification strategy was used. The following primer pairs used were: SMD144 and SMD158 (targeting the 5’-end of *acnB* mRNA); SMD146 and SMD159 (targeting an internal *sdhB* mRNA region); SMD160 and SMD161 (targeting the 5’-end of *gfp* mRNA); SMD162 and SMD163 (targeting the 5’-end of *yfp* mRNA); SMD168 and SMD169 (targeting the 3’-end of *gfp* mRNA); SMD170 and SMD171 (targeting an internal region of *gfp* mRNA). Binding energies were calculated using UNAfold software (Markham *et al.* 2008) and the absence of potential off-targets was checked with RNApredator (Eggenhofer *et al.* 2011). Fragments containing seed sequences complementary to mRNA of the fluorescent markers were transformed into strains containing the respective reporter in the chromosome (KT-GFP or KT-YFP), whilst those with complementarity to *acnB* or *sdhB* transcripts were transformed into *P. putida* KT2440. Electrocompetent *P. putida* KT2440 cells were prepared as described (Choi *et al.* 2006), and electroporated with 500 ng of plasmid DNA. All plasmid constructions were confirmed by sequencing. DNA sequencing and oligonucleotide synthesis were performed by Stab Vida, Portugal.

### Purification of Recombinant <u>P. putida</u> Hfq

*P. putida* KT2440 Hfq protein fused to a histidine tag was overexpressed from a BL21(DE3) *hfq* null strain and purified by affinity chromatography, as described in (Madhushani *et al.* 2015).

### <u>In vitro</u> Transcription

MicC scaffold was generated and ^32^P-labelled by *in vitro* transcription from a synthetic DNA template (MicC_T7, Table S2) with a commercial T7 Promoter similarly to (Milligan *et al.* 1987, Saramago *et al.* 2018). Riboprobe synthesis and oligoprobe labelling were performed as described (Viegas *et al.* 2007). *acnB*, *sdhB* and *micC* scaffold riboprobe templates were generated by PCR with primers SMD122/SMD113, SMD120/SMD114 and MicC_F/MicC_R, respectively.

### Electrophoretic Mobility Shift Assays (EMSA)

^32^P-labelled MicC was diluted in 10 mM of Tris-HCl pH 8 and incubated 10 min at 80°C, followed by 45 min at 37°C to acquire its secondary conformation. 0.075 pmol of MicC were then incubated with increasing concentrations of purified Hfq in Binding Buffer (20 mM Tris-HCl pH 8, 35 mM KCl, 2 mM MgCl_2_), 20 U of RNasin (Promega) and 1 µg of Yeast RNA (Ambion) in volume of 20 μl, at 20°C for 30 min. Binding reactions were mixed with 2μl of loading buffer (48% glycerol, 0.01% bromophenol blue) and electrophoresed on 4% polyacrylamide gel in 0.5X TBE Buffer pH 8 at 200 V. Gels were visualized by PhosphorImaging (FUJI TLA-5100 scanner).

### Spectrofluorimetry Experiments

Total fluorescence was measured along growth in M9 medium using 96-well plates in a FLUOstar OPTIMA (BMG Labtech). Background was determined against control *P. putida* KT2440 carrying pSEVA238-MicC plasmid. Overnight cultures were diluted to OD_600_~0.05 and induced with 1 mM or 5 mM of 3-methylbenzoate (3-MB). Samples were measured in triplicates (200 μl). Wavelengths were set to 485 nm and 520 nm for both reporters, GFP and YFP. The gain was set to 1000 for GFP and to 1750 for YFP. Total fluorescence was calculated as described (de Jong *et al.* 2010). The results are a mean of at least 5 independent experiments.

### Enzymatic Activities

Overnight cultures were diluted to OD_600_~0.05 and induced with 1 mM of 3-MB for 150 min. Aconitase activity was measured as before (Tavares *et al.* 2011) by following the production of NADPH (⃇_340 nm_ = 6.22 mM cm^−1^). Succinate dehydrogenase activity was measured as before (Fernandes *et al.* 2001), using the phenazine methosulfate (PMS)-coupled reduction of dichlorophenolindophenol (DCPIP) (⃇_578 nm_ = 18 mM^−1^cm^−1^). One unit was defined as the enzyme catalysing conversion of 1 nmol of substrate per minute, at 25°C, under the experimental conditions used.

### RNA Extraction and Northern Blotting

Overnight cultures were diluted and immediately induced with 1 mM of 3-MB, incubated 150 min, or grown to a cell density of 0.5 at OD_600_ before induction. Total RNA was extracted using Trizol reagent (Ambion). 15 µg of total RNA was then separated under denaturing conditions either by 8.3M urea/8% polyacrylamide gel in TBE or by 1% agarose MOPS/formaldehyde gel, and Northern blot performed as described (Viegas *et al.* 2007). Visualization was performed in FUJI TLA-5100 scanner and analysis with ImageQuant software (GE Healthcare).

## Results

### Construction of a synthetic sRNA-based expression system

Most synthetic sRNAs are engineered by fusing a target specific sequence to a naturally occurring sRNA scaffold. In a previous work aimed at designing synthetic sRNAs for metabolic engineering in *E. coli*, a screen of 101 *E. coli* sRNAs has selected the MicC scaffold for its superior repression capability (Na *et al.* 2013). Thus, although a MicC homologous sRNA could not be found in *P. putida*, we wanted to test if MicC could be used as a scaffold in this bacterium. Since recruiting of Hfq to the sRNA scaffold should stabilize the sRNA and facilitate its hybridization with the target mRNA as well as mRNA degradation (Aiba 2007), binding of Hfq is an important step in the sRNAs-mediated regulation of gene expression. Thus, we first wanted to test if *P. putida* Hfq would recognize and bind the scaffold of E. coli MicC. In order to do that, we have overexpressed and purified *P. putida* Hfq to near homogeneity (Supplementary Fig. S1A) and tested its ability to bind the MicC scaffold by EMSA (Electrophoretic Mobility Shift Assay). As shown in Supplementary Fig. S1B, incubation of free MicC scaffold with increasing concentrations of purified Hfq led to the formation of Hfq-MicC complexes, which were observed concomitantly with the disappearance of free MicC. This result confirms the ability of *P. putida* Hfq to bind the *E. coli* MicC scaffold and further validates its utilization as a synthetic sRNA scaffold in *P. putida*. Therefore, in order to construct an inducible synthetic sRNA expression system functional in *P. putida*, the *E. coli* MicC scaffold was cloned into the pSEVA238 plasmid (Fig. 1A) yielding pSEVA238-MicC. In this plasmid, gene expression is driven from the inducible *P. putida* native promoter XylS/*Pm*. This regulator/promoter system has been widely used for regulated gene expression and it is suitable for use in metabolic engineering (recently reviewed in (Gawin *et al.* 2017). MicC scaffold was cloned exactly in the +1 position relative to the *Pm* promoter, so that addition of a target specific sequence (seed sequence) immediately upstream of MicC scaffold will allow the expression of a target specific synthetic sRNA after induction (Fig. 1B).

**Figure 1.**
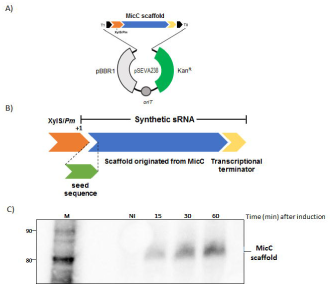
pSEVA238-based synthetic sRNAs expression system. **A)** Insertion of MicC scaffold into pSEVA238 generated a functional segment which includes the XylS/*Pm* promoter (in orange), the MicC scaffold (in blue) with its transcriptional terminator (in yellow). Constructs were assembled in the pSEVA238 backbone vector bearing a kanamycin marker (Kan^R^), the pBBR1 origin of replication, the terminators T_1_ and T_0_ and the origin of transfer *oriT*. **B)** Expression of synthetic sRNAs from *P. putida* XylS/*Pm* promoter. Each construct is composed of a seed sequence (in green), inserted upstream the scaffold of MicC (in blue) in the +1 position of the XylS/*Pm* promoter (in orange). The MicC transcriptional terminator is shown in yellow. **C)** Expression of MicC scaffold after induction of XylS/*Pm*. Northern blot analysis of RNA samples extracted at the time points indicated on top of the image, after induction of XylS/*Pm* with 0.5 mM of 3MB. 15 μg of RNA (each lane) were separated on an 8% polyacrylamide/8.3 M urea gel. The gel was then blotted to a Hybond-N+ membrane and hybridized with a specific probe for MicC sRNA. Details of RNA extraction and Northern blot procedure are described in Materials and Methods. The radiolabelled Decade Marker RNA (Ambion) is on the left side. The respective sizes are represented in nucleotides (MicC scaffold is 79 nts long).

Before proceeding, it was of interest to check out MicC scaffold expression from this system. To this end, Northern blot experiments were conducted and MicC scaffold expression was evaluated before and after induction of KT2440 carrying pSEVA238-MicC with 3-methylbenzoate (3MB). No MicC expression was detected before induction. However, a band corresponding to MicC scaffold could be readily detected 15 min after 3MB addition and an increased expression could be observed 60 min after induction (Fig. 1C). Taken together these results substantiate the utilization of this system for the expression of MicC-based synthetic sRNAs in *P. putida* and encourage its further utilization for modulation of gene expression in this bacterium.

### Downregulation of <u>P. putida</u> cellular targets

The usefulness of this system is dependent on whether it will fit applications on the regulation of cellular endogenous genes. Thus, downregulation of gene expression by the pSEVA238-MicC regulatory system was first assayed in the context of endogenous cellular genes. Since enzymes of the TCA cycle may be preferential targets for redesigning *P. putida* metabolism, aconitase B and succinate dehydrogenase B were the selected candidates in this work. Two sequences complementary to *acnB* and *sdhB* (genes encoding aconitase B and succinate dehydrogenase B enzymes, respectively) mRNAs were designed and introduced into pSEVA238-MicC upstream of the MicC scaffold. We wanted to test sRNA interaction with different regions of the target mRNA molecules. Thus, the sequence chosen for *acnB* targetting hybridized with its 5’-UTR, encompassing the RBS and extended to the translation start codon, whilst a sequence complementary to the coding region was chosen for *sdhB* (Fig. 2A). The outcome was inspected by determining the aconitase or succinate dehydrogenase activities in cellular extracts after respectively inducing the expression of the sRNA directed to *acnB* or *sdhB* targets. As a control, the activity of the same strain expressing the MicC scaffold without the seed sequences was measured in the same conditions. Our data showed that in cells of *P. putida* expressing the sRNAs directed to *acnB* and *sdhB* transcripts, the aconitase and succinate dehydrogenase activities decreased by approximately 40% and 20%, respectively, relative to control sample cells (Fig. 2B). These results indicate that the expression of the specific synthetic sRNAs led to lower protein levels, as judged by the decreased enzyme activities determined in the cell extracts.

**Figure 2.**
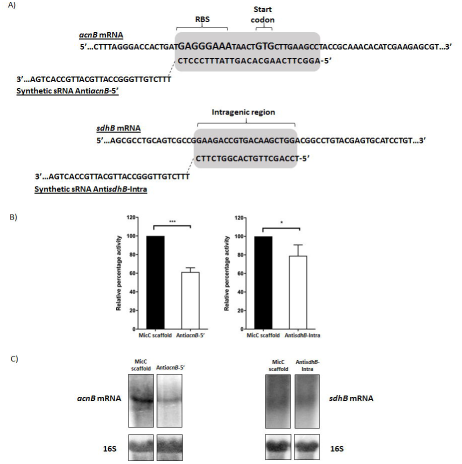
Downregulation of TCA enzymes following expression of specific synthetic sRNAs. **A)** Predicted hybridization regions of the synthetic sRNAs with the TCA cellular targets. The sequences of the hybridization regions of the sRNAs with the respective targets are represented. For simplicity, only a small region of the target genes sequences is shown (dots are used to represent the remaining sequences). Anti*acnB*-5’ hybridizes in the 5’-region of the target, covering the RBS and the translation initiation codon (GTG). Anti*sdhB*-intra hybridizes in the target coding sequence. **B)** Enzymatic activities of aconitase (AcnB) and succinate dehydrogenase (SdhB) measured in extracts of *P. putida* expressing the synthetic sRNAs Anti*acnB*-5’ or Anti*sdhB*-intra, respectively. As control (black bars) enzyme activities were measured on cells expressing only the MicC scaffold. Enzyme activities were measured in cell extracts of *P. putida* induced with 1 mM 3MB for 150 min. Error bars represent the mean of two independent determinations done in duplicate. Asterisks represent statistically significant data relative to the control cells. *P < 0.05, ***P < 0.0001. **C)** Expression of *acnB* and *sdhB* mRNAs after the respective induction of Anti*acnB*-5’ and Anti*sdhB*-intra as stated on top of the images. The control lanes (MicC Scaffold) show the expression of the same transcripts after induction of MicC scaffold. Northern blot analysis of RNA samples extracted 150 min after induction of XylS/*Pm* with 1 mM 3MB. 15 μg of RNA (each lane) were separated on an 1% agarose gel. The gel was then blotted to a Hybond-N+ membrane and hybridized with specific probes.

The reduction of the protein levels could be due to changes in the amount of the target mRNAs, derived from direct mRNA destabilization by the synthetic sRNAs or inhibition of translation, or both. We decided to evaluate the levels of the target mRNAs to discriminate between these hypotheses. The expression levels of the synthetic sRNAs and respective target mRNAs was analysed by Northern blot. As expected, expression of the synthetic sRNAs Anti*acnB*-5’ and Anti*sdhB*-intra was readily observed after induction of XylS/*Pm* with 3MB (Supplementary Fig. S2). The concomitant decreased level of the *acnB* mRNA observed after 3MB induction is indicative of direct destabilization of this message by the sRNA Anti*acnB*-5’ (Fig. 2C). The same was not observed when analysing the *sdhB* message, as no changes could be detected in its amount before or after Anti*sdhB*-intra induction, indicating no evidence of altered mRNA stability (Fig. 2C). Thus, in this case only differences in translation efficiency might be accounting for the final protein level.

Together, these results demonstrate the ability of MicC-based synthetic sRNAs to regulate the level of cellular mRNAs in *P. putida*, whether by interfering with the degradation mechanism or with the translation of the target mRNAs or possibly acting simultaneously at both levels.

### Modulation of reporter genes expression

As a proof of concept, it was of interest to try this system on the regulation of reporter genes. Modulation of reporter genes expression by the pSEVA238-MicC regulatory system was assayed on strains individually carrying the reporter genes GFPmut3 and eYFP (Table S1). We have designed sequences complementary to the translation initiation region (TIR) of each reporter gene in order to create specific synthetic sRNAs directed to the 5’-end of the reporter transcripts. Again, hybridization of the seed sequence with the respective mRNA is expected to change its expression level. To experimentally validate this hypothesis, the fluorescence level of the reporters was evaluated by spectrofluorimetry after induction of the respective synthetic sRNAs. As control, the fluorescence level of the same strains containing only the MicC scaffold (without the seed sequence) was measured in the same conditions. For both reporters, a reduced level of fluorescence was detected after expression of the synthetic sRNAs containing the seed sequences, in comparison with the fluorescence level of the respective control strains (Fig. 3A). According to these results, the fluorescence signal emitted by GFP and YFP was lowered by about 25% and 20%, respectively. Thus, as observed for the cellular targets, expression of these synthetic sRNAs might be impairing translation and/or destabilizing the target mRNAs, leading to a decreased level of the respective fluorescent protein, and ultimately decreasing the fluorescence emitted by the strains. This confirms our previous results with the endogenous cellular genes and further validates this system as a promising toolbox to modulate gene expression in *P. putida*.

**Figure 3.**
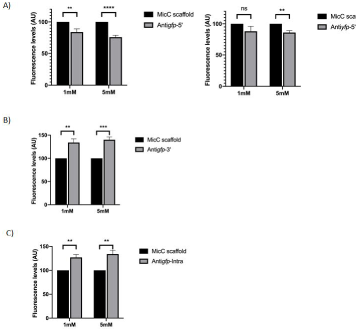
Modulation of reporter genes expression by specific synthetic sRNAs. Fluorescence was measured in growing cultures 150 min after induction of XylS/*Pm* with 1 or 5 mM of 3MB. Total fluorescence was measured in a FLUOstar OPTIMA machine (BMG Labtech). Fluorescence values in arbitrary units (AU) were corrected to the values of the negative control (*P. putida* KT2440 carrying pSEVA238-MicC) and represented as percentage of the basal fluorescence of the control strain (the same strain, carrying the respective reporter and pSEVA238-MicC). Error bars represent the mean of at least five independent measurements. Asterisks represent statistically significant data relative to the control cells. Ns, not significant, *P > 0.05, **P ≤ 0.01, ****P ≤ 0.0001. A) Fluorescence emission of *P. putida* cultures expressing GFP (on the left) or YFP (on the right) after induction of sRNAs directed to the 5’-end of the respective mRNAs (grey bars) or MicC scaffold (black bars) as control. B) Fluorescence emission of *P. putida* cells expressing GFP after induction of a synthetic sRNA directed to the 3’-end of the transcript (grey bar) or MicC scaffold (black bar) as control. C) Fluorescence emission of *P. putida* cells expressing GFP after induction of a synthetic sRNA directed to the coding sequence of the transcript (grey bar) or MicC scaffold (black bar) as control.

sRNAs are potent modulators of gene expression, with the versatile ability not only to downregulate the expression of their targets, but also with the aptitude to stabilize and/or increase translation of the targeted transcripts, as illustrated by a handful of examples in the literature (reviewed by (Saramago *et al.* 2014)). This versatility, which is related with the hybridization region, may be especially interesting in this context, since it should be possible to apply the same synthetic sRNA system to downregulate or upregulate a gene of interest. The choice of the region where the sRNA will hybridise with the target is determinant for the type of control, and only by changing the seed complementarity to a different region of the transcript, it should be possible to differently modulate its expression. As such, we thought that by directing the seed sequence to a different region of the *gfp* transcript that would change its expression differently. This hypothesis was addressed by designing a sequence complementary to the 3’-end of the *gfp* mRNA, a region generally known to have a great impact on RNA stability by being targeted by several exoribonucleases (that cleave RNA in the 3’-5’ direction). It was also of interest to test what would be the changes in the GFP expression level caused by a sRNA directed to an internal region of its transcript. For that, complementary sequences to those regions were introduced as seed sequences, upstream of the MicC scaffold and the alterations in the GFP expression level were again evaluated by spectrofluorimetry. In agreement with our expectations, the fluorescence level of the strain expressing the synthetic sRNA directed to the 3’-end of the transcript was raised by about 40% (Fig 3B). In Fig 3C, we also observed a fluorescence increment after induction of the synthetic sRNA that targets an internal region of the *gfp* transcript (~35%). Although we cannot exclude effects at the translation level, mRNA stabilization is a more plausible effect when the 3’-region was targeted, due to the preferred action of exoribonucleases on that region. The higher GFP expression observed after sRNA hybridization to the internal *gfp* mRNA region might also be due to some stabilizing effect due to occlusion of an endoribonuclease recognition site.

Together these results demonstrate that the type of regulation (up- or down-regulation) achieved for a given gene is determined by the choice of the mRNA region targeted by the synthetic sRNA. Therefore, this is a versatile system for modulation of gene expression, with potential to be used for different purposes in synthetic biology.

## Discussion

In this work we describe the construction and application of a genetic tool for the modulation of gene expression in *P. putida*. Due to the polluted habitats where this robust bacterium thrives, it is equipped with a large number of naturally evolved biochemical and physiological traits that makes it a highly attractive production system for industrial needs (Nikel *et al.* 2018). For this reason, the development of such a regulatory system for specific application in metabolic engineering in this host is of paramount importance.

This genetic tool relies on the induced expression of synthetic sRNAs specifically directed to a chosen target gene. Despite its specificity, we have managed to show the wide applicability of this tool that can be used to modulate the expression of virtually any given gene. The choice of the targeted gene is done by simply altering the “seed sequence”, a small sequence (20-30 nts) that by being complementary to the mRNA target, changes the specificity of the regulatory sRNA. But the utmost advantage of this system is its versatility, due to its aptitude not only to downregulate but also to overexpress any gene of interest. This dual system may thus be used to plug-in and -out genetic circuits for different proposes in synthetic biology.

For the construction of this genetic tool we have used an inducible plasmid from the SEVA collection, in which the expression of the synthetic sRNA is driven from the well-known *P. putida* promoter XylS/*Pm*. Induction of this promoter can be graded by using different, non-expensive, benzoic acid derivatives, which enter cells by passive diffusion and operate in a dose-dependent manner (reviewed by (Gawin *et al.* 2017)). It combines a high expression level with a low fraction of non-induced cells (Calero *et al.* 2016). Cloning of the *E. coli* MicC scaffold under the control of this promoter allowed to detect its expression readily 15 min after induction with 3MB, whereas no expression was detected in the non-induced samples, confirming the usability of the XylS/*Pm* promoter in this system. Further validation of the chosen sRNA scaffold came after binding-shift results, which confirmed the ability of *P. putida* Hfq to recognize and bind the *E. coli* MicC scaffold. Indeed, our results clearly demonstrate the ability of the MicC-based synthetic sRNAs to regulate the expression of several targets in *P. putida*.

sRNAs are potent regulators of gene expression and, although there is a higher number of examples of target downregulation, a few cases of target upregulation have also been described (Saramago *et al.* 2014). The target sequence, the mRNA secondary structure adopted, and the antisense interaction region are among the important factors for the final outcome. In this work we have thus decided to test various interactions regions, comprising the whole mRNA molecules of the genes under study. sRNA hybridization in the 5’-region of the mRNA, comprising the RBS and the start codon, is often associated with translational inhibition due to blockage of ribosome access (Sharma *et al.* 2007, Bouvier *et al.* 2008, Guillier *et al.* 2008, Prevost *et al.* 2011); for review see (Papenfort *et al.* 2009, Saramago *et al.* 2014). This antisense association region is thereby the preferred one when aiming to decrease target expression. In agreement with this expectation, the sRNAs directed to the 5’-end of the target mRNAs – Anti*acnB*-5’, Anti*gfp*-5’ and Anti*yfp*-5’ - resulted in decreased expression levels. This is, however, not always the case of the sRNAs targeting the 5’-end. There are even some examples of sRNAs promoting translation by relieving a hairpin that blocks the RBS upon binding (Soper *et al.* 2010, Saramago *et al.* 2014). Indeed, we have tested two others synthetic sRNAs directed to distinct regions in the 5’-end of the *acnB* and *sdhB* transcripts that caused no changes in the respective enzymatic activities. Despite they were directed to the 5’-end of both mRNAs, the translation process seems not to be compromised (data not shown). Alterations in mRNA turnover caused by the sRNAs may counteract or synergistically favor the translational effects and the final outcome is always the cumulative result of both mechanisms. Accordingly, analysis of the *acnB* mRNA by Northern blot also revealed decreased mRNA levels after Anti*acnB*-5’ expression, which is indicative of mRNA destabilization by the sRNA. The double stranded RNA (dsRNA) region created by the sRNA-target mRNA hybridization may generate a specific cleavage site for the dsRNA specific endoribonuclease RNase III (Viegas *et al.* 2011), presumably leading to the reduced *acnB* transcript level. Changes in the mRNA structure caused by the sRNA hybridization are also known to generate new cleavage sites for other ribonucleases, namely RNase E, an endoribonuclease that specifically recognizes and cleaves single-stranded RNA (ssRNA) (Viegas *et al.* 2008, Storz *et al.* 2011, Bandyra *et al.* 2012). Irrespective of the ribonuclease acting, both effects (inhibition of translation and mRNA destabilization) are probably accounting for the lower aconitase activity obtained upon Anti*acnB*-5’ sRNA induction. Lower succinate dehydrogenase activity was also observed when expressing the synthetic Anti*sdhB*-intra sRNA. In this case however, only translational effects might be occurring, and mRNA destabilization does not appear to be determinant for the final measured activity levels, as no difference was observed in the *sdhB* mRNA levels. The reduced translation might be due to changes in the mRNA structure triggered upon sRNA binding, which would hinder the ribosome access to the transcription start site. Interestingly targeting an *sdhB* intragenic region resulted in decreased enzyme activity, whereas targeting a *gfp* intragenic region resulted in increased fluorescence. Despite the discrepancy, this example illustrates once again the myriad of details when dealing with sRNA regulation, depending on the particularity of each sRNA-target mRNA pair (Villa *et al.* 2018). We may speculate that in the case of the Anti*gfp*-intra sRNA, the antisense hybridization may be affecting mRNA turnover by blocking an endoribonuclease cleavage site, ultimately leading to higher *gfp* mRNA levels. Hypothetically, translation may be promoted through alterations in the mRNA structure following sRNA interaction, or both situations may occur simultaneously accounting for the final fluorescence levels measured. Induction of the sRNA directed to the 3’-end (Anti*gfp*-3’) also raised the fluorescence levels. The most probable reason for this result is stabilization of the target mRNA by limiting the access of the exoribonucleases to the 3’-end of the *gfp* transcript. Two exoribonucleases that specifically degrade RNA in the 3’ to 5’ direction are present in *P. putida* – RNase R and PNPase (Favaro *et al.* 2003, Arraiano *et al.* 2010).

sRNAs are known to regulate complex networks through antisense interaction with their targets, and we have managed to change the expression of several endogenous and reporter genes by using specific synthetic sRNAs. Our work also emphasises the dynamics and complexity of this process. However, this complexity is what gives rise to the great potential of this regulatory system, presumably allowing not only different repression but also overexpression efficiencies. Like in an “haute-couture” atelier, by carefully defining the regions to be targeted, this genetic toolbox allows a tailor-made design for finely tuning the expression of nearly all genes in the cell.

## Acknowledgments

We are grateful to Prof. Victoria Shingler (Umea University, Sweden) for kindly providing us the *E. coli* BL21(DE3)-*Δhfq::cat*-S4 strain. We also acknowledge Maximilian Ole Bahls, from Prof. Sven Panke Lab (ETH, Switzerland), for kindly providing KT-GFP and KT-YFP strains. We are thankful to Dr. Esteban Martinez from Prof. Victor de Lorenzo Lab (CNB, Spain) for supplying pSEVA238 plasmid. We also thank Isabel Pacheco and Teresa Baptista da Silva for technical assistance.

