## Supplementary Material for "Tailor-made sRNAs: a toolbox to control metabolic targets"

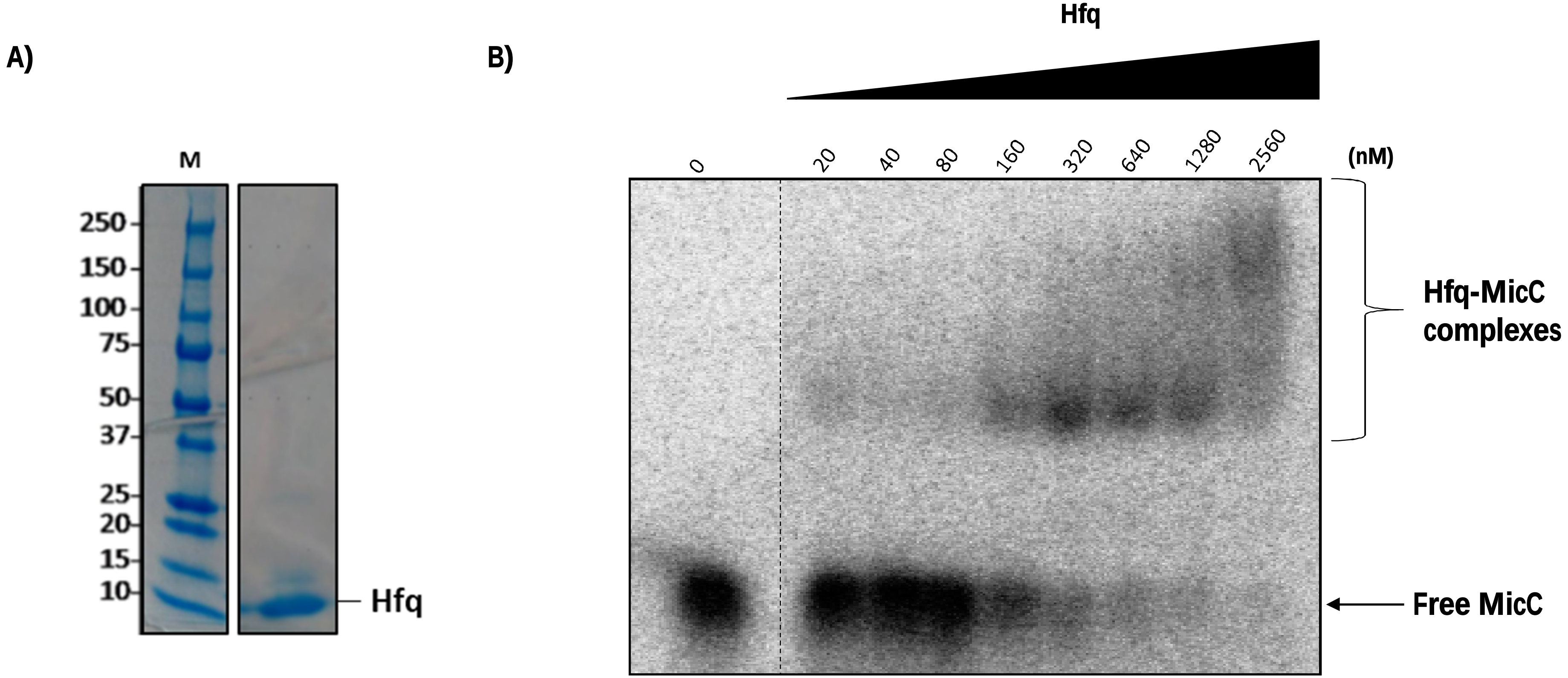


**Figure S1** – **Purified *P. putida* KT2440 Hfq binds *E. coli* MicC Scaffold**. **A)** Purified *P. putida* Hfq (~10.2 kDa) was separated on a 4-15% gradient polyacrylamide gel (Bio-Rad) and visualized by BlueSafe staining. M – Molecular weight marker (Precision Plus Protein Prestained Standards – Bio-Rad) is shown on the left side of the image. **B)** Analysis of the complex formation between the *E. coli* MicC scaffold and the *P. putida* Hfq chaperone by EMSA.

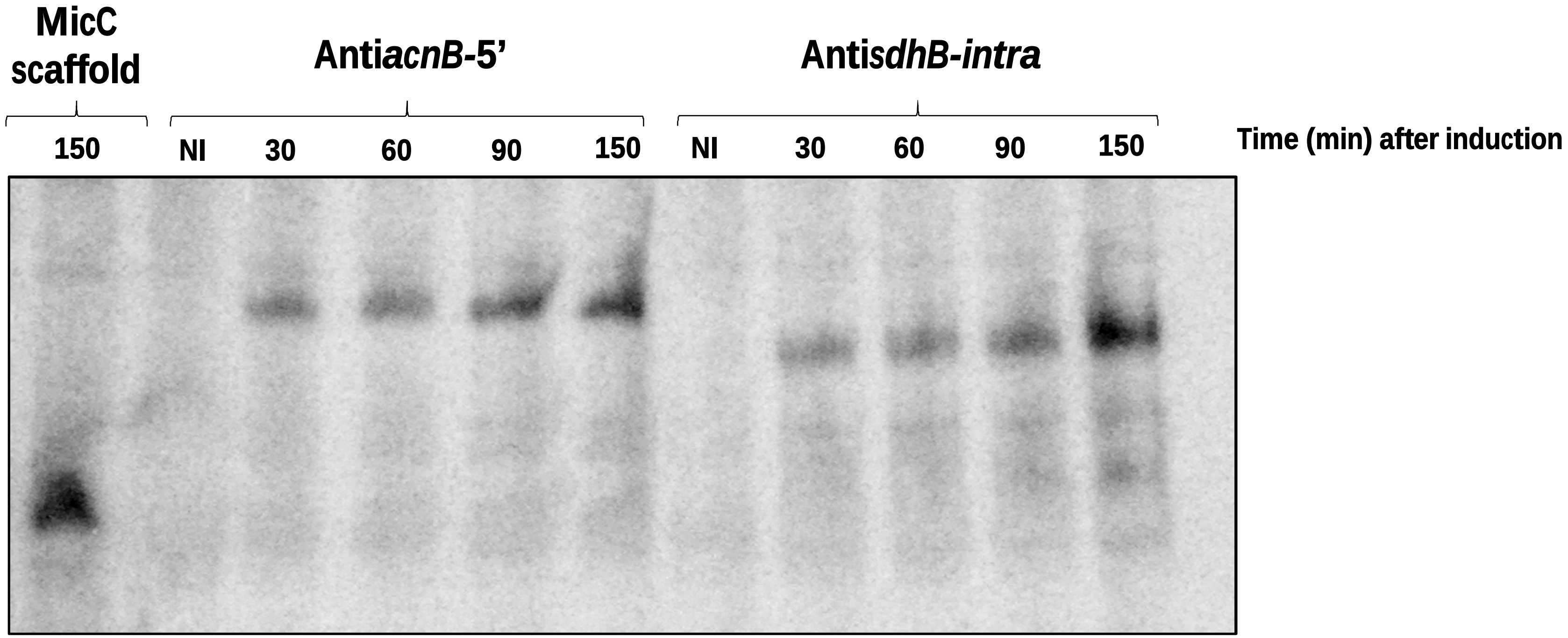


**Figure S2 - Expression of MicC sRNA scaffold, Anti*acnB*-5' and Anti*sdhB*-intra after induction of XylS/Pm promoter.** Northern blot analysis of RNA samples extracted at the time points indicated on top of the image.

**Table S1 –**Strains and plasmids used in this work.

| Strain/Plasmid | Relevant characteristics | Reference/Source |
|---|---|---|
| **Bacteria** | | |
| ***E. coli*** | | |
| BL21(DE3)- Δ*hfq*::*cat*-S4 | $P_{T7}$-*PP hfq-His;* $Cb^R$ expression plasmid (host for Hfq expression and purification) | (Madhushani *et al.* 2015) |
| ***P. putida*** | | |
| KT2440 | mt-2 derivative cured of the TOL plasmid pWW0 | (Bagdasarian *et al.* 1981) |
| KT-GFP | KT2440 with constitutive fluorescent marker (GFP) | (Espeso *et al.* 2016) |
| KT-YFP | KT2440 with constitutive fluorescent marker (YFP) | Victor de Lorenzo's lab* |
| CMA616 | KT2440 carrying pSEVA238-MicC | This work |
| CMA617 | KT2440 carrying pSEVA238- Anti*acnB*-5' | This work |
| CMA618 | KT2440 carrying pSEVA238-Anti*sdhB*-intra | This work |
| CMA619 | KT-GFP carrying pSEVA238-MicC | This work |
| CMA620 | KT-GFP carrying pSEVA238-Anti*gfp*-5' | This work |
| CMA621 | KT-YFP carrying pSEVA238-MicC | This work |
| CMA622 | KT-YFP carrying pSEVA238-Anti*yfp*-5' | This work |
| CMA623 | KT-GFP carrying pSEVA238-Anti*gfp*-intra | This work |
| CMA624 | KT-GFP carrying pSEVA238-Anti*gfp*-3' | This work |
| **Plasmids** | | |
| pSEVA238 | T0/T1 terminators; XylS/Pm promoter; neomycin phosphotransferase ($Kan^R$) | (Silva-Rocha *et al.* 2013) |
| pSEVA238-MicC | pSEVA238 containing MicC scaffold ($Kan^R$) | This work |
| pSEVA238-Anti*acnB*-5' | pSEVA238-MicC containing Anti*acnB*-5' ($Kan^R$) | This work |
| pSEVA238-Anti*sdhB*-intra | pSEVA238-MicC containing Anti*sdhB*-intra ($Kan^R$) | This work |
| pSEVA238-Anti*gfp*-5' | pSEVA238-MicC containning Anti*gfp*-5' ($Kan^R$) | This work |
| pSEVA238-Anti*yfp*-5' | pSEVA238-MicC containing Anti*yfp*-5' ($Kan^R$) | This work |
| pSEVA238-Anti*gfp*-intra | pSEVA238-MicC containing Anti*gfp*-intra ($Kan^R$) | This work |
| pSEVA238-Anti*gfp*-3' | pSEVA238-MicC containing Anti*gfp*-3' ($Kan^R$) | This work |

*Victor de Lorenzo Lab., unpublished

**Table S2 -** List of oligonucleotides used in this work.

| Oligo name | Sequence 5' to 3' |
|---|---|
| SMD154 | ATAAAAAGACAAGCCCGAACAGTCGTCCGGGCTTTTTTTACAGAAACAATAATAATGGAGTCATGACC |
| SMD155 | ATGTTGGAAAATCAGTGGCAATGCAATGGCCCAACAGAAATGCATAAAGCCTAAGGGGTAGGCCTTAC |
| SMD144 | AGTTATTTCCCTCTTTCTGTTGGGCCATTGCATTGCCAC |
| SMD158 | GTGCTTGAAGCCTTGCATAAAGCCTAAGGGGTAGGCCTTACTAG |
| SMD146 | CACGGTCTTCTTTCTGTTGGGCCATTGCATTGCCAC |
| SMD159 | ACAAGCTGGATGCATAAAGCCTAAGGGGTAGGCCTTACTAG |
| SMD160 | TCCTTTACGCATTTTCTGTTGGGCCATTGCATTGCCAC |
| SMD161 | GAAGAACTTTTCTGCATAAAGCCTAAGGGGTAGGCCTTACTAG |
| SMD162 | CTTGCTCAGCATTTTCTGTTGGGCCATTGCATTGCCAC |
| SMD163 | GGCGAGGAGCTGTGCATAAAGCCTAAGGGGTAGGCCTTACTAG |
| SMD168 | AATCCCAGCATTTCTGTTGGGCCATTGCATTGCCAC |
| SMD169 | ACACATGGCTTGCATAAAGCCTAAGGGGTAGGCCTTACTAG |
| SMD170 | TTAACAAGGGTTTCTGTTGGGCCATTGCATTGCCAC |
| SMD171 | TAGAATCGAGTGCATAAAGCCTAAGGGGTAGGCCTTACTAG |
| SMD122 | GTTTTTTTTTTAATACGACTCACTATAGGTTTCGCCAGGCACCTTGAAC |
| SMD113 | GGCTACAACATCGAAACGCTGG |
| SMD120 | GTTTTTTTTTTAATACGACTCACTATAGGAGGAACGGCTTCACCTTCTC |
| SMD114 | GGGTGGCTGCTATGTTGAAAGTCG |
| MicC_F | GTTATATGCCTTTATTGTCACAT |
| MicC_R | GTTTTTTTTTTAATACGACTCACTATAGGGAGGCGTTCGGGCTTGTCA |
| MicC_T7 | AAAAAAAGCCCGGACGACTGTTCGGGCTTGTCTTTTTATATGTTGGAAAATCAGTGGCAATGCAATGGC CCAACAGAAATATAGTGAGTCGTATTA |
| T7 promoter | TAATACGACTCACTATA |
| svpa25 | ACGGCTACCTTGTTACGACTT |
